# A Reproducibility Analysis-based Statistical Framework for Residue-Residue Evolutionary Coupling Detection

**DOI:** 10.1101/2021.02.01.429092

**Authors:** Yunda Si, Yi Zhang, Chengfei Yan

**Affiliations:** School of Physics, Huazhong University of Science and Technology, China

**Keywords:** direct coupling analysis, quality control, statistical methods, contact prediction

## Abstract

Direct coupling analysis (DCA) has been widely used to infer evolutionary coupled residue pairs from the multiple sequence alignment (MSA) of homologous sequences. However, effectively selecting residue pairs with significant evolutionary couplings according to the result of DCA is a non-trivial task. In this study, we developed a general statistical framework for significant evolutionary coupling detection, referred to as IDR-DCA, which is based on reproducibility analysis of the coupling scores obtained from DCA on manually created MSA replicates. IDR-DCA was applied to select residue pairs for contact prediction for monomeric proteins, protein-protein interactions and monomeric RNAs, in which three different versions of DCA were applied. We demonstrated that with the application of IDR-DCA, the residue pairs selected using a universal threshold always yielded stable performance for contact prediction. Comparing with the application of carefully tuned coupling score cutoffs, IDR-DCA always showed better performance. The robustness of IDR-DCA was also supported through the MSA down-sampling analysis. We further demonstrated the effectiveness of applying constraints obtained from residue pairs selected by IDR-DCA to assist RNA secondary structure prediction.

## Introduction

Contacting residues in monomeric proteins/RNAs or between interacting proteins/RNAs often show covariance in the process of evolution to maintain the architectures and the interactions of these macromolecules, which allows us to infer the intra- or inter-protein/RNA residue-residue contacts through co-evolutionary analysis [1]. Direct coupling analysis (DCA) is a class of widely used methods for co-evolutionary analysis, which quantifies the direct coupling strength between two residue positions of a biological sequence through global statistical inference using maximum entropy models learned from large alignments of homologous sequences [2]. Comparing with local statistical methods like mutual information (MI) and correlated mutation analysis, DCA is able to disentangle direct couplings from indirect transitive correlations, thus showing much better performance in predicting residue-residue contacts [3,4]. A wide variety of algorithms at different levels of approximation for implementing DCA have been developed in recent years, with the focus being on improving the accuracy of DCA and increasing the computational efficiency. The developed algorithms for DCA include message passing DCA (mpDCA) [3], mean-field DCA (mfDCA) [4] and pseudo-likelihood maximization DCA (plmDCA) [2]. Among these developed algorithms, PlmDCA is currently the most popular algorithm for implementing DCA because of its high accuracy and moderate computational cost, which has been successfully applied to directly acquire contact constraints to assist the prediction of protein/RNA structures, interactions and dynamics [5–14], or been used to provide major feature components for deep learning-based contact/distance prediction methods [15–19].

Comparing with so many efforts made on improving DCA algorithms and applying DCA to obtain structural information from sequence data, relatively less attention has been made on how to quantify the number of residue pairs with significant evolutionary couplings and select the predictive residue pairs from the result of DCA. Generally, residue pairs with higher coupling scores obtained from DCA tend to have higher probabilities to form contacts. Therefore, in most previous studies, often a certain number (e.g. top 10 or top L/5, L as the sequence length) of residue pairs with the highest coupling scores were selected for contact prediction, or a coupling score cutoff was set empirically to select residue pairs with coupling scores higher than the cutoff. However, both the number of predictive residue pairs and the coupling score values are influenced by many factors including the number and the length of the homologous sequences forming the MSA, the detailed settings of the DCA algorithm, the functional characteristics of the macromolecule [6,7]. Therefore, neither applying a “number” cutoff nor a “coupling score” cutoff is an ideal protocol for selecting predictive residue pairs from the result of DCA. In a previous work, for predicting residue-residue contacts between interacting proteins, Ovchinnikov et al. first rescaled the raw coupling scores from Gremlin (a software for implementing plmDCA) with an empirical model to consider the influence of the number and the length of the homologous sequences forming the MSA on the coupling scores, then determined an optimal score cutoff based on the inter-protein residue-residue contacts in the crystal structure of the 50S ribosome complex [6]. For the same purpose, Hopf *et al.* did something similar but rescaled the coupling scores from EVcouplings (another software for implementing plmDCA) with a different empirical model [7]. Both the two methods achieve success in selecting inter-protein residue pairs for contact prediction. However, since the parameters of these empirical models were tuned on only a limited number of cases, whether they are applicable for more general cases is questionable. Besides, Xu et al. proposed a statistical approach referred to as inverse finite-size scaling (IFSS) to estimate the significance of DCA results, which was later applied in epistasis detection of microbial genomes [20–22]. However, to the best of our knowledge, the effectiveness of this approach has never been shown in selecting evolutionary coupled residue pairs. The lack of a general approach for detecting significant evolutionary couplings from the result of DCA limits the appropriate application of this method. For example, when applying DCA to infer inter-protein residue pairs with significant evolutionary couplings to assist the protein-protein interaction prediction at large scale, without appropriately measuring the coupling significance, we may introduce false positive couplings or miss significant couplings.

In this study, we develop a general statistical framework for significant residue-residue coupling detection. The development of this statistical framework is inspired by the quality control protocols in functional genomic experiments, in which often reproducible signals in multiple experimental replicates are considered as the genuine functional signal [23,24]. Here, given an MSA of homologous sequences, two MSA (pseudo) replicates are created by randomly assigning the sequences in the MSA into two groups. DCA is then performed on both the original full MSA and the two MSA replicates. We assume that the significant couplings are reproducible from DCA on the two MSA replicates. Therefore, we perform reproducibility analysis on the coupling scores obtained from DCA on the two MSA replicates, from which we assign each residue pair an irreproducible discovery rate (IDR) calculated with the Gaussian copula mixture modelling described in Li *et al.* [25], with the lower the IDR, the more reproducible the residue-residue coupling. Then, we create an IDR signal profile for the residue pairs under consideration, which represents the IDR variation with the ranking of the residue pairs sorted descendingly according to the coupling scores obtained from DCA on the full MSA. The residue pairs before the IDR signal profile reaching a certain threshold are considered to be with significant evolutionary couplings. This statistical framework, referred to as IDR-DCA, was applied to select residue pairs for contact prediction for 150 monomeric proteins, 30 protein-protein interactions and 36 monomeric RNAs, in which the DCA were performed with three different versions of DCA including EVcouplings [26], Gremlin [5] and CCMpred [27]. The result shows that IDR-DCA can effectively select evolutionary coupled residue pairs with a universal threshold (IDR cutoff=0.1), for that the numbers of residue pairs selected by IDR-DCA vary dramatically for cases across the three datasets, but the accuracies of the selected residue pairs for contact prediction are kept stable. Comparing with the application of the DCA tool specific coupling score cutoffs carefully tuned on each dataset to reproduce the overall accuracies of the residue pairs selected by IDR-DCA, IDR-DCA is always able to select more residue pairs, and provide effective contact predictions for more cases. We further evaluated the robustness of IDR-DCA through the MSA downsampling analysis. The result shows that as the numbers of homologous sequences forming the MSAs getting smaller and smaller, generally IDR-DCA would select fewer and fewer residue pairs to keep the accuracy of the selection, but the advantage of IDR-DCA over the application of coupling score cutoffs are always kept at different levels of the MSA down-sampling. Therefore, IDR-DCA provides an effective and robust statistical framework for selecting evolutionary coupled residue pairs.

## Results

### 1. Overview of IDR-DCA

IDR-DCA includes three major stages: creating pseudo-replicates, performing reproducibility analysis and detecting significant couplings, which are described in detail in the following subsections (See Figure 1).

**Figure 1.**
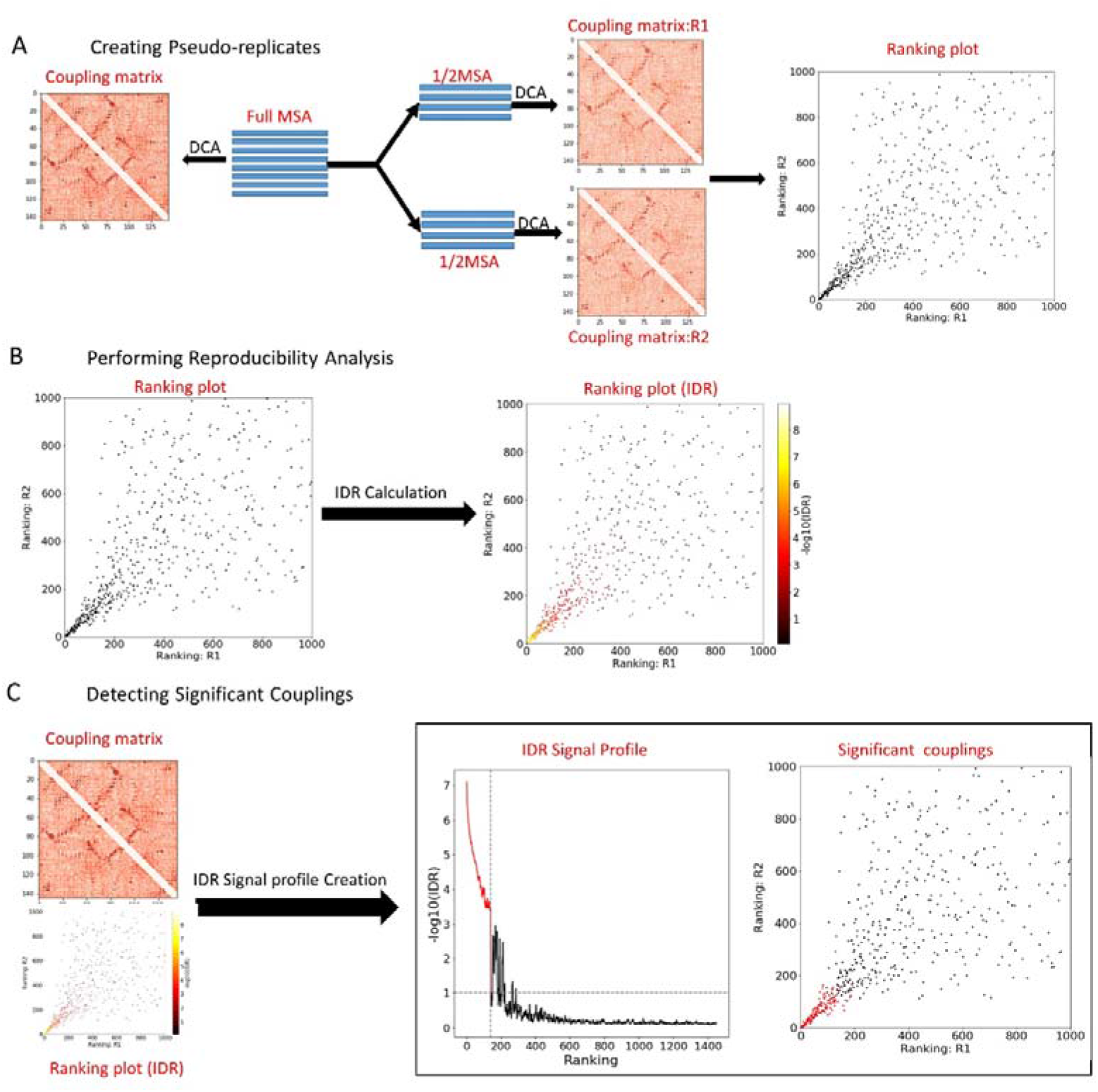
The flowchart of IDR-DCA. (A) Creating pseudo MSA replicates for DCA; (B) Performing reproducibility analysis; (C) Detecting significant couplings.

#### 1.1 Creating pseudo-replicates

As it is shown in Figure 1A, given an MSA of homologous sequences, we first perform DCA on the full MSA, from which we can obtain a coupling score *x_i_* for each residue pair. The residue pairs are then sorted descendingly rankings *n_i_* are more likely to be with significant evolutionary couplings. After according to the coupling scores, in which the residue pairs with higher that, the aligned sequences in the MSA are randomly grouped into two subsets separately, from which we can obtain a coupling score tuple (*x_i,1_,x_i,2_*) and a without realignment, and we then perform DCA on the two MSA subsets ranking tuple (*n_i,1_,n_i,2_*) for each residue pair, with *x_i,1_, x_i,2_* representing the coupling scores for the residue pair *i* from the DCA on the two MSA subsets, and *n_i,1_, n_i,2_* representing the rankings of residue pair *i* sorted according to the coupling scores descendingly. Since the two MSA subsets can be considered as (pseudo) biological replicates, the significant couplings are expected to be reproducible from the DCA on the two MSA replicates. Therefore, we perform reproducibility analysis on the coupling scores obtained from the replicated DCA to evaluate the reproducibility of each residue-residue coupling. It should be noted that if the provided MSA contains a large number of redundant sequences (including extremely similar sequences), insignificant couplings may also show a certain level of reproducibility. To avoid reproducible couplings caused by the issue of sequence redundancy, redundant sequences in the MSA should be filtered.

#### 1.2 Performing reproducibility analysis

Since the scale and the distribution of the coupling scores obtained from DCA are case dependent, it is not appropriate to measure the reproducibility of the residue-residue coupling through the direct comparison of the coupling score values from the two MSA replicates. In this study, we measure the reproducibility of each residue-residue coupling through calculating the irreproducible discovery rate (IDR) for each residue-residue coupling with the Gaussian copula mixture modelling described in Li *et al* [25], in which the rankings rather than the coupling score values from the two MSA replicates were employed in the statistical modeling. Specifically, we assume that there are two types of residue pairs (i.e. evolutionary coupled residue pairs and evolutionary uncoupled residue pairs), for which the observed coupling score tuples (*x_i,1_,x_i,2_*) are generated by a latent variable tuple (unobserved) (*z_1_, z_2_*) following the Gaussian mixture distribution (*π_0_h_0_(z_1_,z_2_)+π_1_h_1_(z_1_,z_2_)*), with

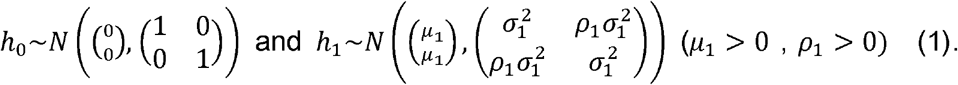

Where *h_0_* and *h_1_* correspond to the uncoupling component and the coupling component respectively, and *π_0_* and *π_1_* are the corresponding weights of the two components. Since evolutionary coupled residue pairs generally have higher and more reproducible coupling scores, we require (*μ*_1_ > 0, *ρ_1_* > 0). Because (*z_1_*, *z_2_*) are not observable, and the relationship between (*z_1_*, *z_2_*) and the observable coupling score tuples (*x_i,1_,x_i,2_*) is unknown, the association parameters 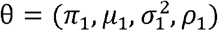 of the Gaussian mixture model are determined through maximizing the likelihood function of corresponding copula mixture model:

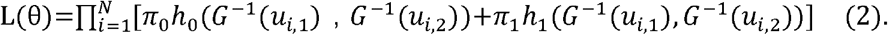

Where 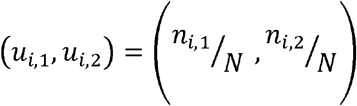 is the normalized ranking tuple of residue pair *i*, with *n_i,1_,n_i,2_* corresponding to the rankings of residue pair *i* according to the coupling scores from replicated DCA, and N representing the total number of residue pairs; 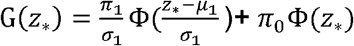 is the cumulative marginal distribution of *z_1_* and *z_2_*, with *z* representing either *z_1_* or *z_2_*, and Φ representing the standard normal cumulative distribution function. As we can see from Equation (2) that only the rankings obtained from the two MSA replicates are employed in the parameter determination.

Given a set of parameters θ, the probability that a residue pair with the normalized ranking tuple (*u_i,1_,u_i,2_*) being an evolutionary uncoupled residue pair (local IDR) can be computed as:

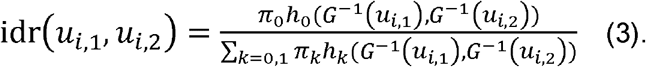

The local IDR are then converted to (global) IDR for the multiple hypothesis correction. The IDR of each residue pair represents the reproducibility of the corresponding residue-residue coupling, with the lower the IDR value, the higher the reproducibility (See Figure 1B).

#### 1.3. Detecting significant couplings

The rankings (*n_i_*) and the reproducibilities (IDRs) of residue-residue couplings are unified for the significant coupling detection. Specifically, we build an IDR signal profile for all residue pairs under consideration, which represents the IDR variation with the ranking of the residue pairs sorted descendingly according the coupling scores obtained from DCA on the full MSA. Generally, the IDR of each residue-residue coupling increases (-log10(IDR)↓) with fluctuation when its ranking goes down. After smoothing the IDR signal profile using a moving average filter with a window size 5, the residue pairs before the IDR signal reaching a specified cutoff are considered to be with significant evolutionary couplings (See Figure 1C).

### 2. Detecting significant evolutionary couplings with variable IDR cutoffs

IDR-DCA was used to detect intra-protein residue-residue couplings for the 150 monomeric proteins in the original PSICOV contact prediction dataset [28], inter-protein residue-residue couplings for 30 protein-protein interactions from Ovchinnikov *el al.* [6], and intra-RNA residue-residue couplings for 36 monomeric RNAs from Pucci *et al.* [13], in which the DCA were performed with three widely used plmDCA-based DCA software including EVcouplings [26], Gremlin [5] and CCMpred [27]. We first applied variable IDR cutoffs to select evolutionary coupled residue pairs. The percentage of contacting residue pairs in the selected residue pairs was used to evaluate the accuracy of the selection. Two intra-protein residues were considered to be in contact if their Cβ-Cβ distance (Cα-Cα distance in the case of glycine) is smaller than 8Å. For the inter-protein residues, the distance cutoff was relaxed to 12 Å considering that the inter-protein residues have much lower contact probability than the intra-protein residues. For the intra-RNA residues, a contact was defined if their C1′-C1′ distance is smaller than 12 Å. In Figure 2A-2C, we show the overall accuracies of the selected residue pairs from each dataset with the application of variable IDR cutoffs. As we can see from Figure 2A-2C that independent on the DCA tools, for all the three datasets, the accuracies drop at a relative slow speed when increasing IDR cutoff until reaching 0.1, and after that the accuracies drop dramatically. Therefore, 0.1 can be considered as a natural IDR cutoff for selecting residue pairs for contact prediction when using the IDR-DCA statistical framework.

**Figure 2.**
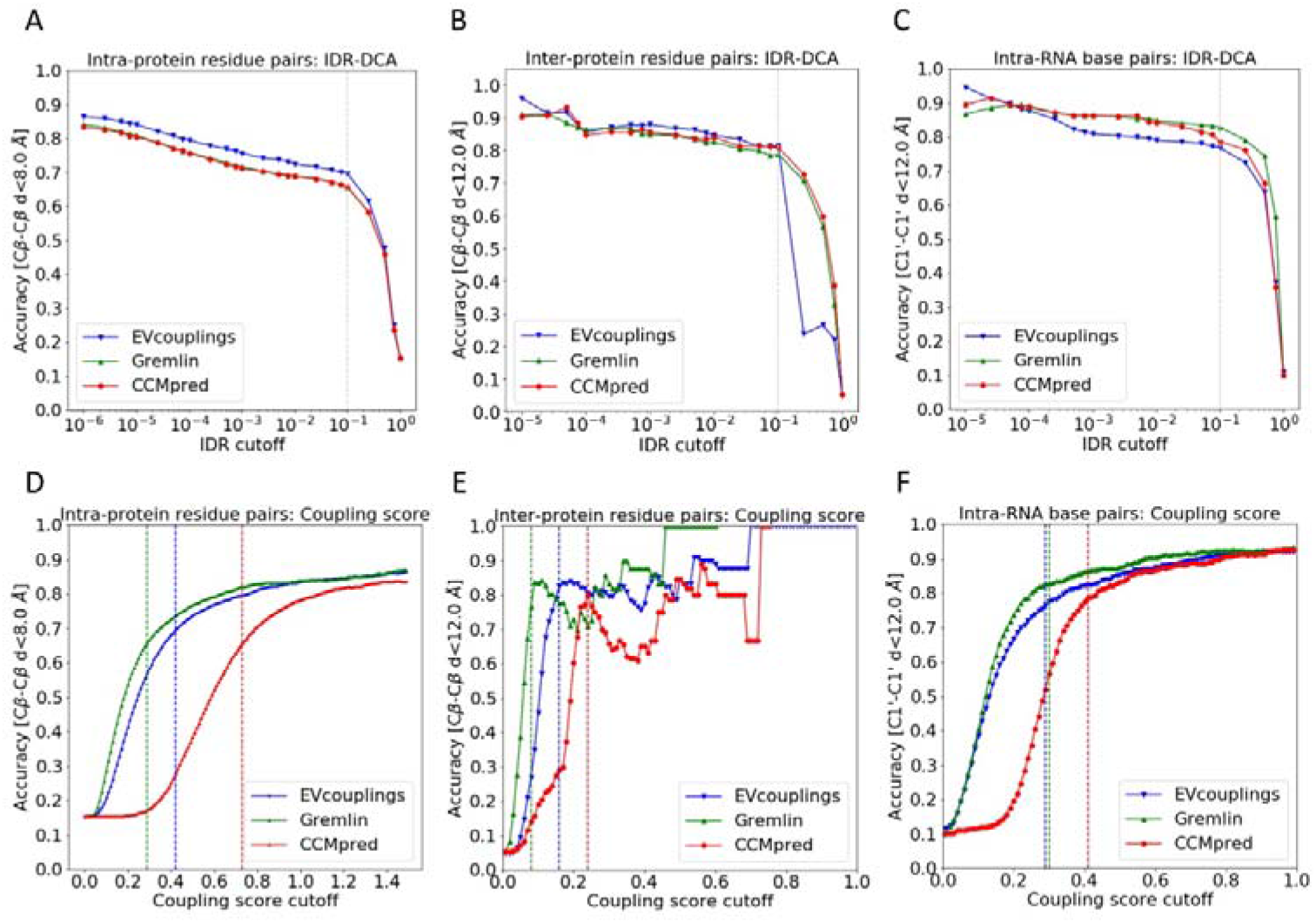
The overall accuracies of residue pairs selected from each dataset based on IDR-DCA and coupling scores with the application of variable IDR and coupling score cutoffs. (A)-(C) The overall accuracies of residue pairs selected by IDR-DCA with the application of variable IDR cutoffs from the three datasets: (A) The monomeric protein dataset; (B) The protein-protein interaction dataset; (C) The monomeric RNA dataset. (D)-(F) The overall accuracies of residue pairs selected based on coupling scores with the application of variable coupling score cutoffs from the three datasets: (D) The monomeric protein dataset; (E) The protein-protein interaction dataset; (F) The monomeric RNA dataset. EVcouplings, Gremlin and CCMpred were applied to perform the DCA for each case in the three datasets respectively. The grey vertical dashed lines in (A)-(C) represent the natural IDR cutoff (0.1) for IDR-DCA. The blue, green and red vertical dashed lines in (D)-(F) represent the empirical coupling score cutoffs for EVcouplngs, Gremlin and CCMpred respectively, which were tuned on each dataset to reproduce the accuracies of residue pairs selected by IDR-DCA with 0.1 as the IDR cutoff.

For the purpose of comparison, we also selected residue pairs based on the coupling score values. In Figure 2D-2F, we show the overall accuracies of the residue pairs selected from each dataset with the application of variable coupling score cutoffs. As we can see from Figure 2D-2F, the accuracy of the selected residue pairs varies with the choice of the coupling score cutoff in DCA tool specific and dataset dependent ways. Therefore, a universal coupling score cutoff is not applicable for selecting residue pairs for contact prediction. For each dataset, we can set empirical DCA tool specific coupling score cutoffs for the residue pair selection, with which the selected residue pairs reproduce the accuracies of the residue pairs selected by IDR-DCA with 0.1 as the IDR cutoff. Specifically, for the monomeric protein dataset, the coupling score cutoffs for EVcouplings, Gremlin and CCMpred were set as 0.42, 0.29 and 0.73; for the protein-protein interaction dataset, the coupling score cutoffs were set as 0.16, 0.09 and 0.24; and for the monomeric RNA dataset, the coupling score cutoffs were set as 0.29, 0.30 and 0.41. The dramatic variations of the coupling scores cutoffs between different DCA tools and different datasets show that the coupling score is not a good metric for the predictive residue pair selection.

It is easy to explain that the obtained coupling score cutoffs are tool dependent. Since all the three versions of DCA use some sort of regularizations to avoid the model overfitting, if the parameters of the regularizations are set differently, or the regularizations are done in different ways, the scales of the obtained coupling score values will vary. This is also the reason that the standard practice in the DCA application relies more on the order of the prediction than numeric values of the coupling scores. Besides, it is also reasonable that different datasets show different scales of coupling scores, since the different types of biophysical interactions can have different strengths. For example, the intra-protein residue-residue interactions are generally stronger and more conserved than the inter-protein residue-residue interactions, and may also show different evolutionary characteristics from the RNA residue-residue interactions.

### 3. The performance of IDR-DCA with a universal IDR cutoff (0.1) on evolutionary coupled residue pair selection

We analyzed the performance of IDR-DCA on selecting evolutionary coupled residue pairs with 0.1 as the IDR cutoff. In Figure 3A-3C, we show the accuracies, the numbers and the corresponding coupling score cutoffs (i.e. the smallest coupling score) of the selected residue pairs for each case in the three datasets. As we can see from Figure 3A-3C, the selected residue pairs yield quite stable accuracies across cases in the three datasets independent on the DCA tools (e.g. for most of the cases, the accuracies of selected residue pairs are higher than 50%.), although the numbers of the selected residue pairs vary dramatically. We can also see that the corresponding coupling score cutoffs of the selected residue pairs vary dramatically not only between DCA tools, but also across cases in the three datasets. This further supports that it is not appropriate to apply a universal coupling score cutoff to select residue pairs for contact prediction.

**Figure 3.**
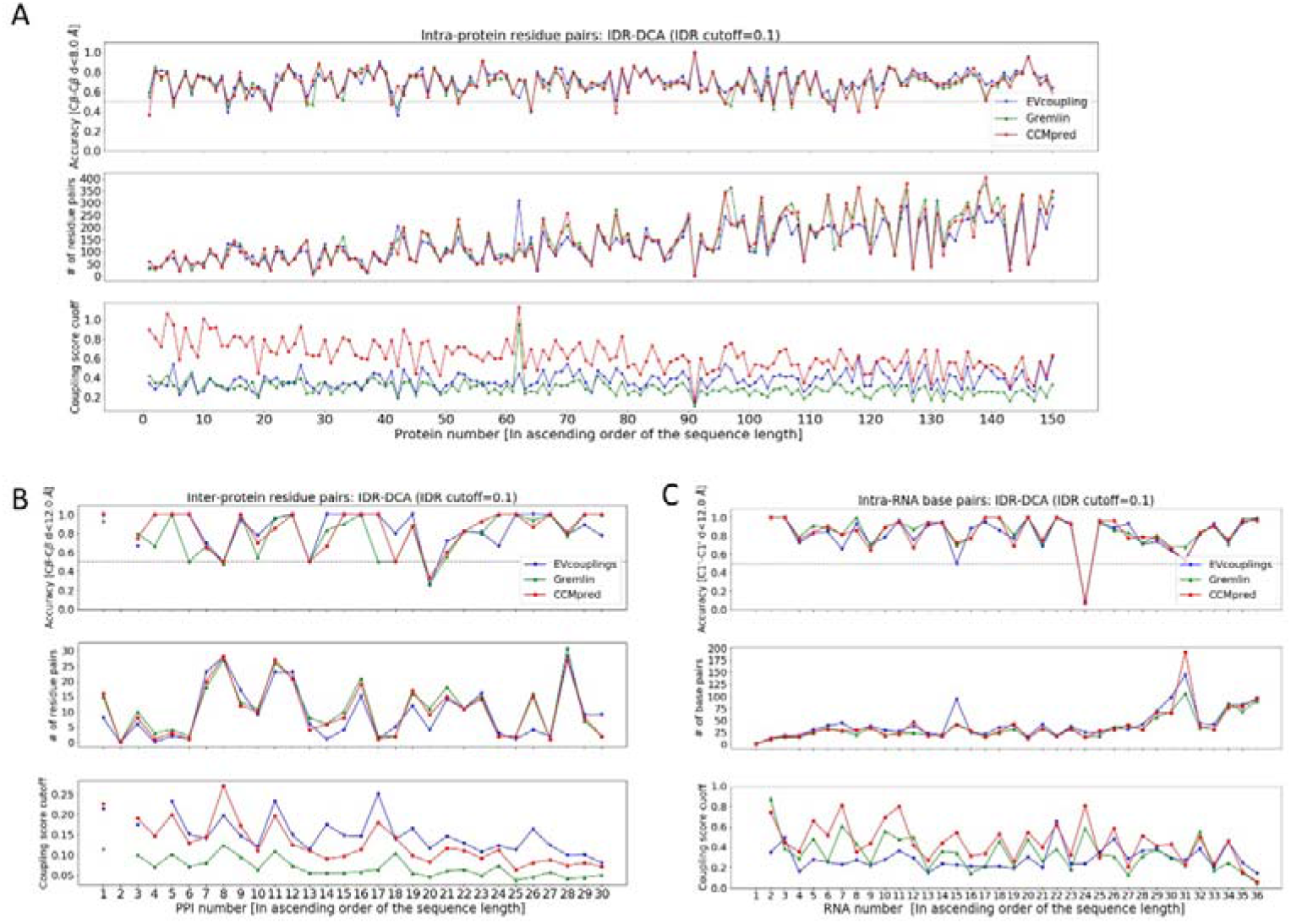
The performance of IDR-DCA on evolutionary coupled residue pair selection with 0.1 as the IDR cutoff. (A)~(C) The accuracies, the numbers and the corresponding coupling score cutoffs of the residue pairs selected by IDR-DCA with 0.1 as the IDR cutoff for each case in the three datasets: (A) The monomeric protein dataset; (B) The protein-protein interaction dataset; (C) The monomeric RNA dataset. For each case, EVcouplings, Gremlin and CCMpred were applied to perform the DCA respectively. In the case that no residue pair is selected by IDR-DCA, the corresponding accuracy and the corresponding score cutoff is not shown. For each dataset, the cases (proteins, protein-protein interactions, RNAs) are ordered ascendingly in the plot according to their sequence lengths.

We further analyzed the distance distribution of non-contacting intra-protein residue pairs (Cβ-Cβ distance ≥8Å) selected by IDR-DCA from the monomeric protein dataset. The analysis was focused on the intra-protein residue pairs for which have the largest sample size. We found that the distances of most of the non-contacting residue pairs selected by IDR-DCA are just slightly larger than 8Å (e.g. <12Å), as it is shown in Figure S1A. Therefore, we suspect that many of these “non-contacting” residue pairs by definition may also be “truly” evolutionary coupled. We also noticed a tiny fraction of the selected residue pairs are in long distance (e.g. ≥12Å in the crystal structure. These residue pairs can be evolutionary coupled with the long distances caused by protein conformational changes, as it is shown by Anishchenko et al., in which they found that most of the evolutionary coupled residue pairs not in repeat proteins are actually in spatial proximality in at least one biologically relevant conformation [11]. Besides, the alignment errors in MSA and the approximations made in DCA can also be responsible for these exceptions.

We also noticed that for several cases in the protein-protein interaction dataset and the monomeric RNA dataset, IDR-DCA was not able to find any evolutionary coupled residue pairs. The scatter plot of the coupling scores of all residue pairs for these cases is shown in Figure S1B, in which the contacting residue pairs are colored red, and the non-contacting residue pairs are colored black. As we can see from the plot, for all the cases, very few top-ranked residue pairs based the coupling scores from DCA are in contacts in the 3D crystal structure. This means that the DCA on these cases failed to correctly model the residue-residue couplings, which might be caused by the lack of effective sequences in their MSAs.

Therefore, it is encouraging that IDR-DCA can avoid selecting false positive residue pairs from these cases. This phenomenon did not happen to our monomeric protein dataset, for the monomeric proteins used in our study are all single domain proteins with large number of homologous sequences in their MSAs, thus the DCA on these cases can always successfully identify a certain number of evolutionary coupled residue pairs.

### 4. The performance comparison between the application of IDR-DCA and coupling score cutoffs on evolutionary coupled residue pair selection

We compared the performance of IDR-DCA on evolutionary coupled residue pair selection with the application of the DCA tool specific coupling score cutoffs tuned on each dataset. As we have described in section 2, the coupling score cutoffs were determined to reproduce the accuracies of the residue pairs selected by IDR-DCA. As we can see from Figure 4A-4C, for all the three datasets, IDR-DCA with a universal IDR cutoff (0.1) is always able to select more residue pairs than the application of the carefully tuned DCA tool specific coupling score cutoffs, although the accuracies of the selected residue pairs are almost the same.

**Figure 4.**
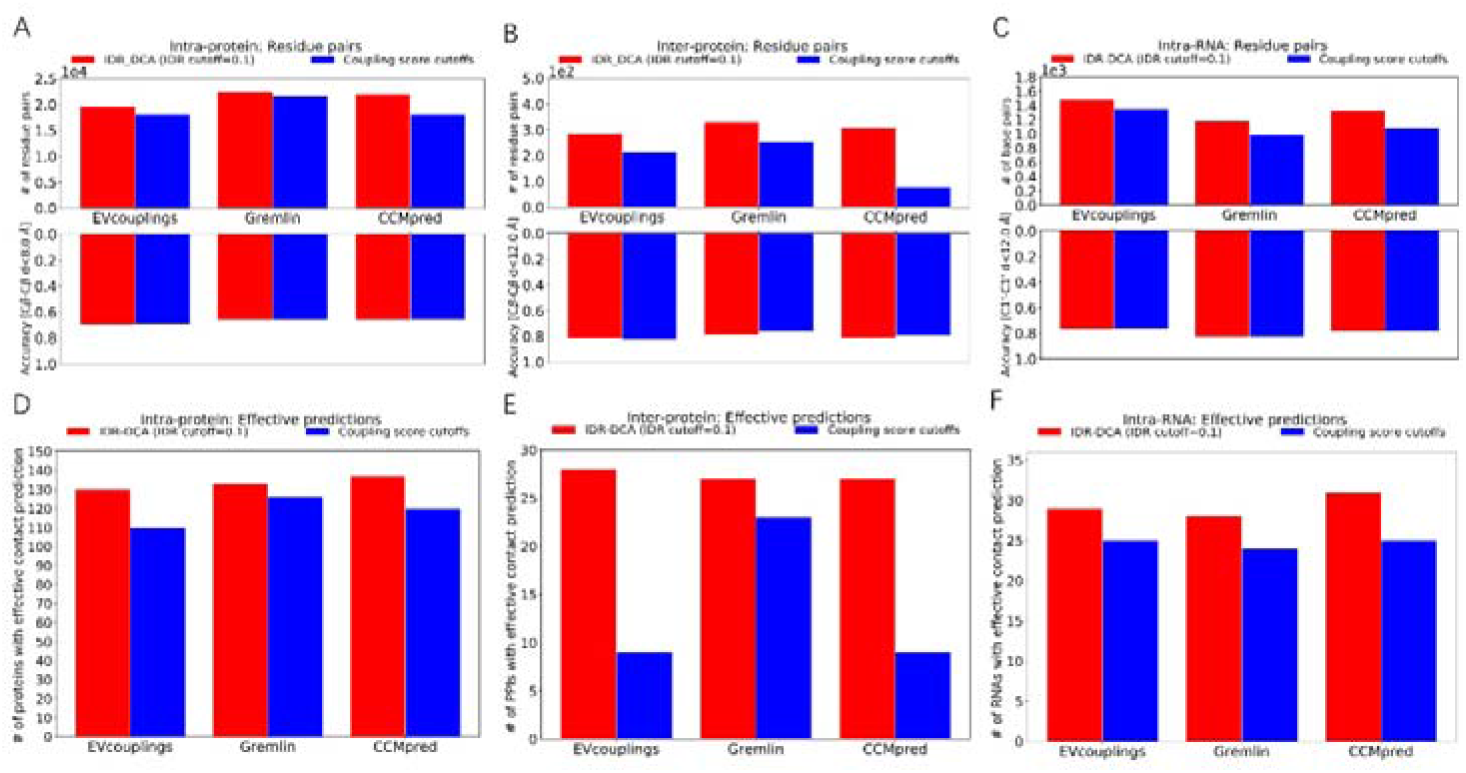
The performance comparison between the application of IDR-DCA (0.1 as the IDR cutoff) and the coupling score cutoffs for the evolutionary coupled residue pair selection. (A)-(C) The comparison of the numbers of the residue pairs selected by applying IDR-DCA with 0.1 as the IDR cutoff and by applying the DCA tool specific coupling score cutoffs from the three datasets: (A) The monomeric protein dataset; (B) The protein-protein interaction dataset; (C) The monomeric RNA dataset. The coupling score cutoffs for the EVcouplings, Gremlin and CCMpred were tuned on each dataset respectively to reproduce the accuracies of residue pairs selected by IDR-DCA with 0.1 as the IDR cutoff. (D)-(F) The comparison of the numbers of cases with effective contact predictions provided by applying IDR-DCA with 0.1 as the IDR cutoff and by applying the DCA tool specific coupling score cutoffs for residue pair selection on the three datasets: (D) The monomeric protein dataset; (E) The protein-protein interaction dataset; (F) The monomeric RNA dataset.

Besides, IDR-DCA also shows a more stable performance across cases in each dataset. For example, for most of the cases, the numbers of residue pairs selected by IDR-DCA are very similar between different DCA tools, but the numbers of residue pairs selected by applying the coupling score cutoffs are highly dependent on the choice of the DCA tools (See Figure S2). Since the differences between the three DCA tools are only on the detailed settings of the plmDCA algorithm (e.g. the initial values for the optimization, the criterion for the convergence, the ways of regularizations, etc.), DCA implemented with the three DCA tools on the same MSA should provide similar number of evolutionary coupled residue pairs.

An effective contact prediction should provide enough residue pairs above a certain level of accuracy to assist the structure prediction. Here, as rules of thumb, for monomeric proteins or RNAs, we defined a prediction providing not fewer than L/5 (L as the sequence length) residue pairs with an accuracy not lower than 50% as an effective contact prediction; for protein-protein interactions, an effective contact prediction was defined if it can provide at least one residue pairs with an accuracy not lower than 50%, considering even one inter-protein residue contact constraint can significantly reduce the configuration space of the protein-protein interactions. It should be noted that the “effective contact prediction” defined here is only to make performance comparison between the application of the universal IDR cutoff and the variable coupling score cutoffs quantitively, thus other reasonable criteria can also be used in the analysis. In Figure 4D-4F, we show the comparison of the numbers of cases with effective contact predictions provided by applying IDR-DCA and by applying the coupling score cutoffs from the three datasets respectively. As we can see that IDR-DCA is always able to provide effective contact predictions for much more cases than the application of the coupling score cutoffs. Besides, we can also see that the performance gap for the protein-protein interaction dataset is much larger than those for the monomeric protein dataset and the monomeric RNA dataset. This is mainly because the coupling scales of DCA methods are also highly dependent on the sequence length. For the monomeric protein and monomeric RNA dataset, the variations of the sequence lengths (51~267 and 24~492) are significantly smaller than that for the protein-protein interaction dataset (181~1453).

### 5. Evaluating the robustness of IDR-DCA through the MSA downsampling analysis

We further evaluated the robustness of IDR-DCA through the MSA down-sampling analysis on the monomeric protein dataset. Specifically, for each protein in monomeric protein dataset, 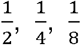 and 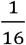 of sequences in the original MSA were randomly selected to form the MSAs with different levels of downsampling, and then we applied IDR-DCA on the downsampled MSAs with 0.1 as the IDR cutoff to select evolutionary coupled residue pairs. EVcouplings, Gremlin and CCMpred were still applied respectively to perform the DCA.

In Figure 5A-5C, we show the accuracies, the numbers of residue pairs selected by IDR-DCA for each case in the monomeric protein dataset with the application of the three tools for DCA respectively. As we can see from the Figure 5A-5C, as the size of MSA getting smaller and smaller, the numbers of the selected residue pairs are also getting smaller and smaller, however, the accuracies of the selected residue pairs for contact prediction are kept stable (See Figure S3 for an example). Since DCA on MSA with fewer sequences tends to have lower statistical power to accurately model the residue-residue couplings, it is reasonable that IDR-DCA selected fewer residue pairs as we kept downsampling the MSA.

**Figure 5.**
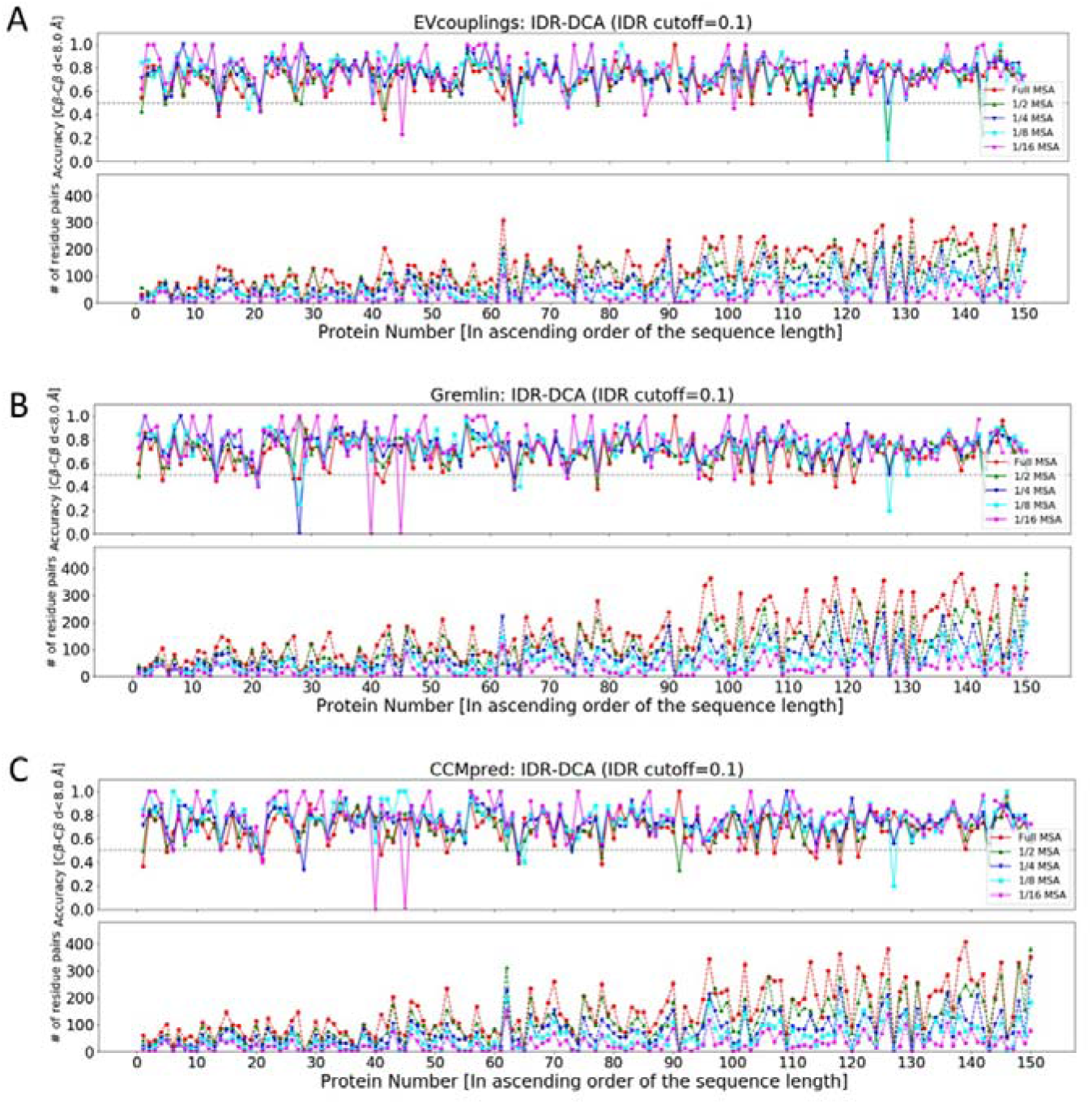
The robustness evaluation of IDR-DCA (0.1 as the IDR cutoff) on evolutionary coupled residue pair selection through the MSA downsampling analysis. (A)-(C) The accuracies and the numbers of the residue pairs selected by IDR-DCA (0.1 as the IDR cutoff) for each protein in the monomeric protein dataset, in which the DCA were performed on the MSAs with different levels of downsampling with the application of the three DCA tools: (A) EVcouplings; (B) Gremlin; (C) CCMpred. In the case that no residue pair is selected, the corresponding accuracy is not shown in the plot. The proteins are ordered ascendingly in each plot according to their sequence lengths.

For the purpose of comparison, we also applied the previous determined coupling score cutoffs to select residue pairs according to the coupling scores obtained from DCA on the MSA with different levels of downsampling (See Figure S4). In Figure 6A-6C, we show the comparison of the numbers and the accuracies of the residue pairs selected by applying IDR-DCA and by applying the coupling scores cutoffs for the three DCA tools respectively. As we can see from Figure 6A-6C, the accuracies of the residue pairs selected by the two approaches are kept comparable across different levels of the MSA downsampling, however, IDR-DCA is always able to select more residue pairs. Besides, we also compared the numbers of proteins with effective contact predictions provided by the two approaches, which is shown in Figure 6D-6F. The definition of an effective contact prediction for the monomeric protein is the same as before. As we can see from Figure 6D-6F, for all the three DCA tools, IDR-DCA is always able to provide effective contact predictions for more proteins across different levels of the MSA downsampling.

**Figure 6.**
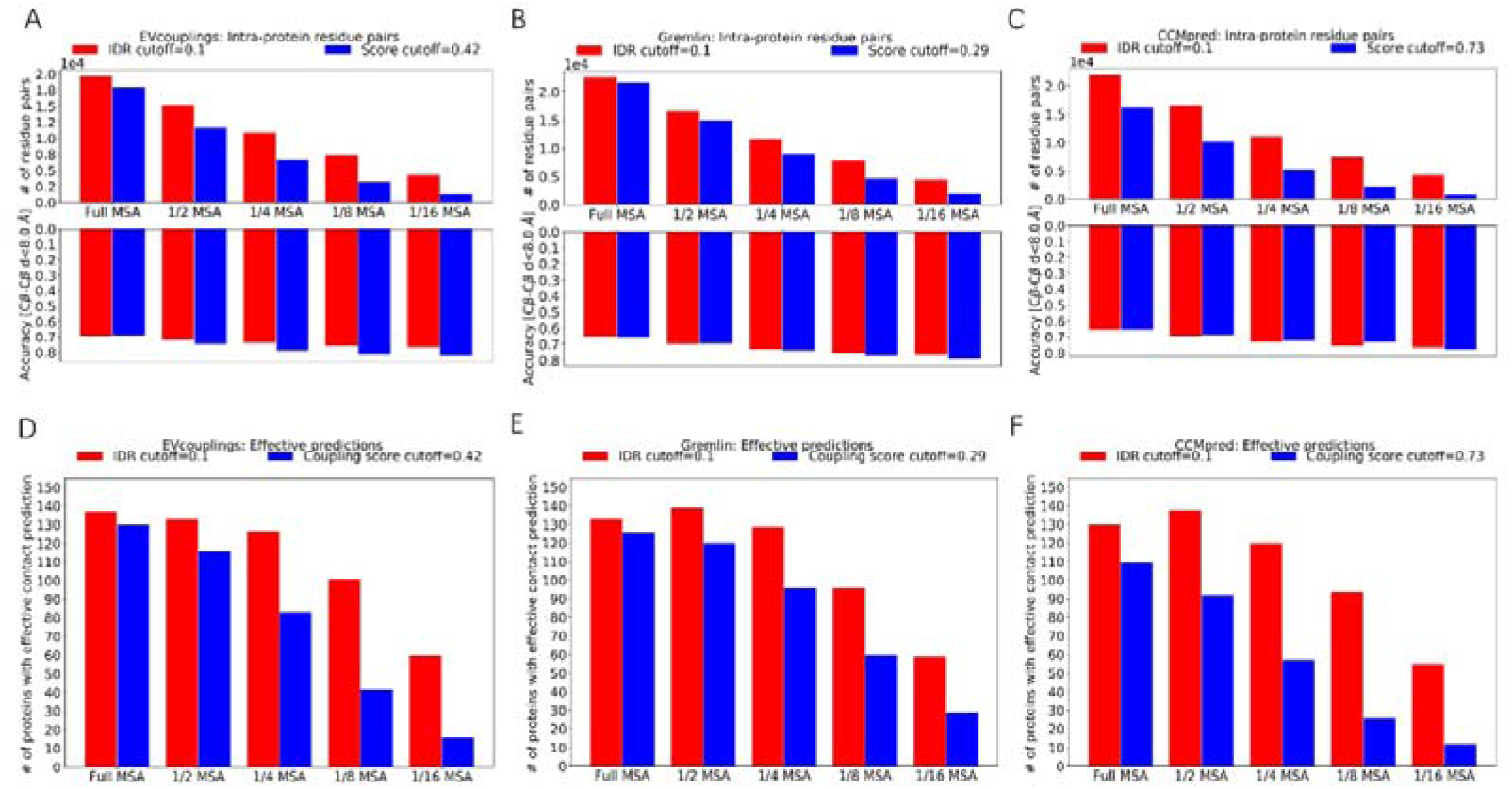
The performance comparison between the application of IDR-DCA (0.1 as the IDR cutoff) and the coupling score cutoffs for the evolutionary coupled residue pair selection from the monomeric protein dataset on the MSAs with different levels of downsampling. (A)-(C) The comparison of the numbers and the accuracies of residue pairs selected by applying IDR-DCA (0.1 as the IDR cutoff) and by applying the coupling score from the monomeric protein dataset, in which the DCA were performed on the MSAs with different levels of downsampling with the three DCA tools: (A) EVcouplings; (B) Gremlin; (C) CCMpred. (D)-(F) The comparison of the numbers of cases with effective contact predictions provided by applying IDR-DCA (0.1 as the IDR cutoff) and by applying the coupling score cutoffs for residue pair selection from the monomeric protein dataset on the MSAs with different levels of downsampling, in which the DCA were performed with the three DCA tools: (D) EVcouplings; (E) Gremlin; (F) CCMpred.

### 6. Applying constraints obtained from IDR-DCA to assist RNA secondary structure prediction

We used RNA secondary structure prediction as an application example to show the benefit of leveraging IDR-DCA statistical framework. Specifically, the webserver 2dRNAdca [29] (http://biophy.hust.edu.cn/new/2dRNAdca/) was applied for the RNA secondary structure prediction, which first applied a remove-and-expand algorithm to refine residue pairs selected by IDR-DCA (0.1 as the IDR cutoff) to form the prior constraints for RNA secondary structure prediction, and then the prior constraints were further used to guide RNAfold [30] (a minimum free energy based RNA secondary structure prediction method) to predict the RNA secondary structure. 26 RNAs without broken strands were selected from the RNA dataset for testing the protocol. Since the result of IDR-DCA is not that dependent on the specific DCA tools, only the IDR-DCA results based on CCMpred were employed in our study. Besides, the prediction performances by RNAfold without prior constraints, with prior constraints refined by the remove-and-expand algorithm from the top L/5 (L as the sequence length) residue pairs and from residue pairs selected according to the coupling scores (CCMpred coupling score cutoff: 0.41) were also evaluated as the references. In Figure 7, we show the Matthews Correlation Coefficients (MCC) between the experimental RNA secondary structure and the predicted secondary structures by the four protocols for each of the 26 RNAs (the RNAs are ordered according to the sequence length ascendingly). As we can see from the figure that the introduction of prior constraints by the three protocols all dramatically improves the prediction performance for most of the cases. However, the prediction protocol using the constraints refined from the residue pairs selected by IDR-DCA yields a more stable performance, especially for large RNAs. We also noticed that for short RNAs (e.g. sequence length<80), the secondary structure prediction with the three types of prior constraints almost makes no difference. This is mainly because that for short RNAs, the number of residue pairs selected by the three approaches are very similar. However, for long RNAs, IDR-DCA can more effectively select the residue pairs with significant evolutionary couplings, thus for which the RNA secondary structure prediction with the IDR-DCA constraints shows better performance (see Table S1). It should be noted that since the variations of the RNA sizes and MSA qualities (the numbers of effective sequences for all the MSAs are larger than 70) of RNA dataset used in our study are not that large, selecting residue pairs according to the coupling scores or trivially selecting the top L/5 residue pairs to some extent can also produce reasonable results. This is also the reason that the predicted RNA secondary structures using the prior constraints from IDR-DCA only achieved slightly higher accuracies. It is reasonable to expect that performance gaps can be enlarged if a more diverse RNA dataset is applied here.

**Figure 7.**
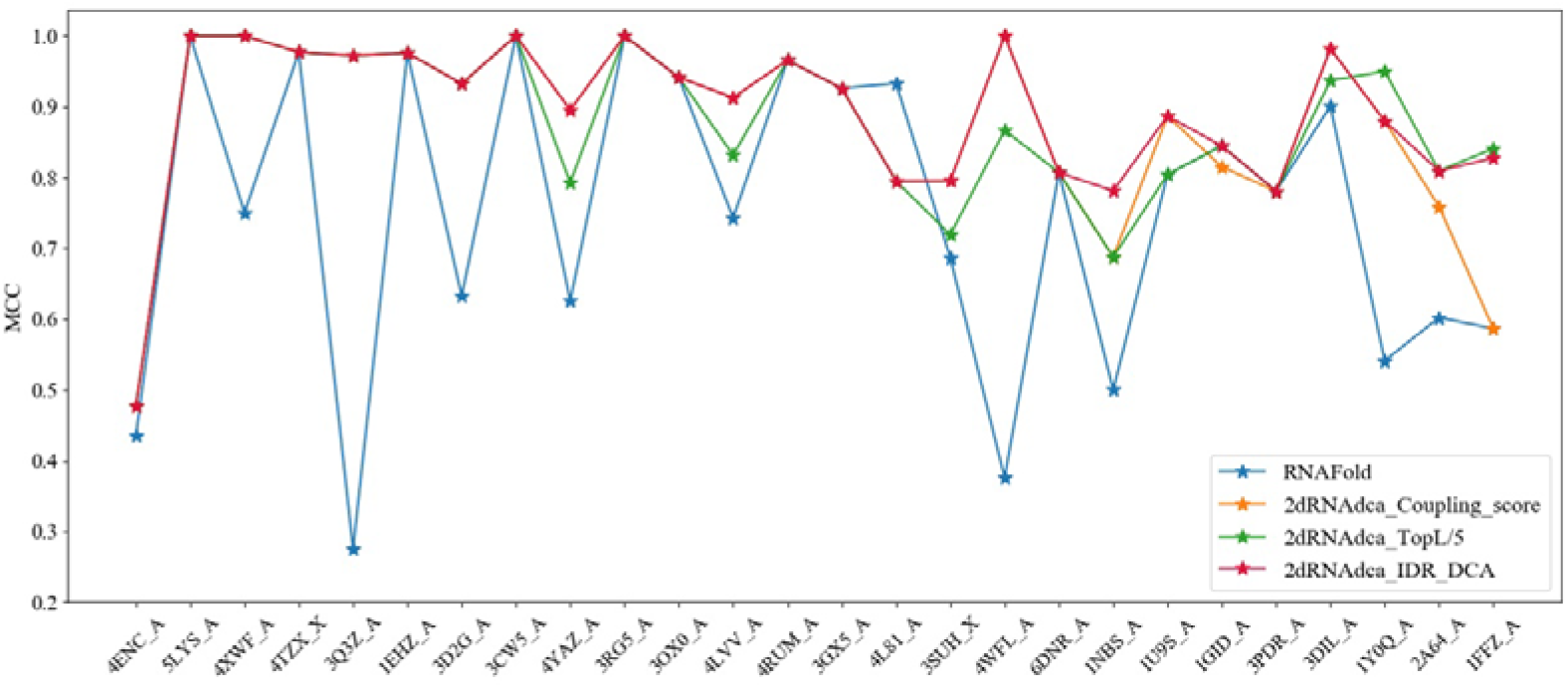
The Matthews Correlation Coefficients (MCC) between the experimental RNA secondary structure and the predicted RNA secondary structures by the four protocols for each of the 26 RNAs. The RNAs are ordered ascendingly according to their sequence lengths.

## Discussion

DCA has been widely used to obtain residue-residue contact information to assist the protein/RNA structure, interaction and dynamics prediction. Besides, the coupling score matrices obtained from DCA also provide major feature components for most of the deep learning methods for the protein residue-residue contact/distance prediction, which has revolutionized the field of protein structure prediction [15]. Given the MSA of homologous sequences, DCA can be easily implemented with the state-of-art DCA software to provide the residue-residue coupling scores. However, it is not easy to quantify the number of residue pairs with significant evolutionary couplings and select these predictive residue pairs from the result of DCA, because the number of predictive residue pairs and the coupling score values from DCA are influenced by many factors including the number and the length of the homologous sequences forming the MSA, the detailed settings of the DCA algorithm, the functional characteristics of the macromolecule, etc.

In this study, we presented a general statistical framework named IDR-DCA for selecting residue pairs with significant evolutionary couplings. Benchmarked on datasets of monomeric proteins, protein-protein interactions and monomeric RNAs, we showed that IDR-DCA can effectively select predictive residue pairs with a universal IDR cutoff (0.1). Comparing with the application of the DCA tool specific coupling score cutoffs carefully tuned on each dataset to reproduce the accuracies of the residue pairs selected by IDR-DCA, IDR-DCA is always able to select more residue pairs and provide effective contact predictions for more cases. Therefore, IDR-DCA provides an effective statistical framework for the evolutionary coupled residue pair detection, which can also be considered as a general approach for controlling the quality of the result of DCA. Besides, we also used RNA secondary structure prediction as application example of IDR-DCA. Of course, IDR-DCA can also be used in other application scenarios. Since the statistical framework of IDR-DCA is not dependent on any detailed implementation of the DCA algorithm, this statistical framework is also expected to be applicable to performing quality control for other data-driven contact prediction methods including deep learning.

## Materials and Methods

### 1. Preparing the three datasets

#### 1.1 The protein dataset

The PSICOV contact prediction dataset [28] which contains 150 proteins was used to evaluate the performance of IDR-DCA on detecting intra-protein residue-residue couplings. The structures of the 150 proteins were obtained from http://bioinfadmin.cs.ucl.ac.uk/downloads/PSICOV/suppdata/. The MSAs of homologous proteins for the 150 proteins were built through searching the whole-genome sequence databases Uniclust30 [31] and UniRef90 [32] and the metagenome database (Metaclust) [33] using DeepMSA [34]. The redundant sequences with sequence identity higher than 90% in the MSA were removed with HHfilter [35].

#### 1.2 The protein-protein interaction dataset

The PDB benchmark from Ovchinnikov *et al.* was used to evaluate the performance of IDR-DCA on detecting inter-protein residue-residue couplings[6].

The complex structures of the 30 protein-protein interactions were downloaded direct from the Protein Data Bank [36]. The MSAs of non-redundant protein-protein interologs for the 30 protein-protein interactions were obtained from the supplementary data of Ovchinnikov *et al* [6]. The original PDB benchmark contains 32 protein-protein interactions, however, 1IXR_B-1IXR_C was removed due to the interacting region of 1IXR_B was missing in the MSA of protein-protein interologs; and 2Y69_B-2Y69_C was removed for the two chains do not directly interact with each other in the crystal structure.

#### 1.3 The RNA dataset

The *D^High^* dataset from Pucci *et al.* [13] containing 36 RNAs associated to RNA families with the number of effective sequences larger than 70 was used to evaluate the performance of IDR-DCA on detecting intra-RNA residue-residue couplings. The structures and MSAs of the 36 RNAs were obtained from https://github.com/KIT-MBS/RNA-dataset. For each MSA, the columns with more than 50% gaps were first removed, and then the redundant sequences with sequence identity higher than 95% were removed with HHfilter.

### 2. Performing the DCA

The three DCA tools: EVcouplings, Gremlin and CCMpred were applied respectively in this study to perform the DCA. Evcouplings (only the plmc module) was obtained from https://github.com/debbiemarkslab/plmc; Gremlin was obtained from https://github.com/sokrypton/GREMLIN; and CCMpred was obtained from https://github.com/soedinglab/CCMpred. CCMpred and Gremlin were run with their default settings, and EVcouplings was run with parameters “-le 16.0 -lh 0.01 -m 100” for proteins and protein-protein interactions, and with parameters “-a .ACGU -le 20.0 -lh 0.01 -m 50” for RNAs according to the recommendations from the website.

### 3. Performing the reproducibility analysis

The R package ‘idr’ obtained from https://cran.r-project.org/web/packages/idr/index.html was employed for the reproducibility analysis with the set of parameters “mu=1.0, sigma=1.0, rho=0.2, p=0.1, eps=1e-5, max.iter=1000”. For the monomeric proteins and RNAs, the residue pairs separated by less than 6 residues were not considered in the IDR estimation. For the protein-protein interactions, considering the contact probability of inter-protein residues is much lower than that of intra-protein residues, the intra- and inter-protein residue pairs were mixed together for the IDR calculation for the purpose of better parametrization of the statistical model. However, only the IDRs of inter-protein residue-residue couplings were used to build the IDR signal profile for the inter-protein evolutionary coupling detection. For the purpose of reducing the computational cost, we only perform the reproducibility analysis for the top 10*L (L as the sequence length) couplings ranked based the coupling score obtained from the DCA on the full MSA, since the number of evolutionary coupled residue pairs is generally much smaller than this value.

### 4. Determining the coupling score cutoffs

For the purpose of comparison, we determined a DCA tool specific coupling score cutoff on each dataset to reproduce the accuracy of the residue pairs selected by IDR-DCA with 0.1 as the IDR cutoff. Specifically, for each DCA tool, starting from 0, we kept increasing the coupling score cutoff for selecting residue pairs from the corresponding dataset with a step size 0.01, until the accuracy of the selected residue pairs exceeded the accuracy of the residue pairs selected by IDR-DCA (0.1 as the IDR cutoff). Then the coupling score cutoff which yielded an accuracy closest to accuracy of IDR-DCA was chosen as the empirical coupling score cutoff for this DCA tool on the corresponding dataset.

### 5. Predicting RNA secondary structure

The 2dRNAdca webserver (http://biophy.hust.edu.cn/new/2dRNAdca/) were employed to perform the constraints assisted RNA secondary structure prediction. Ten RNAs with broken strands were removed from the RNA dataset in the secondary structure prediction. The experimental secondary structure of each RNA was calculated with X3DNA [37] without pseudoknotted base pairs. The predicted secondary structures were evaluated by calculating the Matthews Correlation Coefficient (MCC) between the predicted structure and the experimental structure, which was calculated using the following formula:

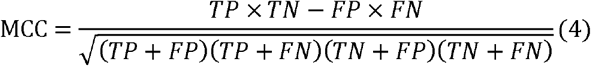

Where TP is the number of true positive base pairs; FP is the number of false positive base pairs; TN is the number of true negative base pairs and FN is the number of false negative base pairs.

#### Key points

- A novel statistical framework is proposed to control the quality of the result of DCA.
- Our method allows to effectively select residue pairs with significant evolutionary couplings using a universal threshold.
- Our method with a universal threshold consistently achieves better performance than carefully tuned coupling score cutoffs.
- Prior constraints obtained from our method has a robust performance in assisting RNA secondary structure prediction.

## Availability

The script for IDR calculation was provided in https://github.com/ChengfeiYan/IDR-DCA.

## Funding

This work is supported by the new faculty startup grant (grant number: 3004012167) of Huazhong University of Science and Technology.

**Yunda Si** is a PhD student in the School of Physics at Huazhong University of Science and Technology. His research interests include protein structure prediction, protein-protein interaction prediction and deep learning.

**Chengfei Yan** is an associate professor in the School of Physics at Huazhong University of Science and Technology. His research interests include molecular docking, protein-protein interaction prediction and biological data mining.

