## Supplemental Figures for "A Reproducibility Analysis-based Statistical Framework for Residue-Residue Evolutionary Coupling Detection"

### Supplementary

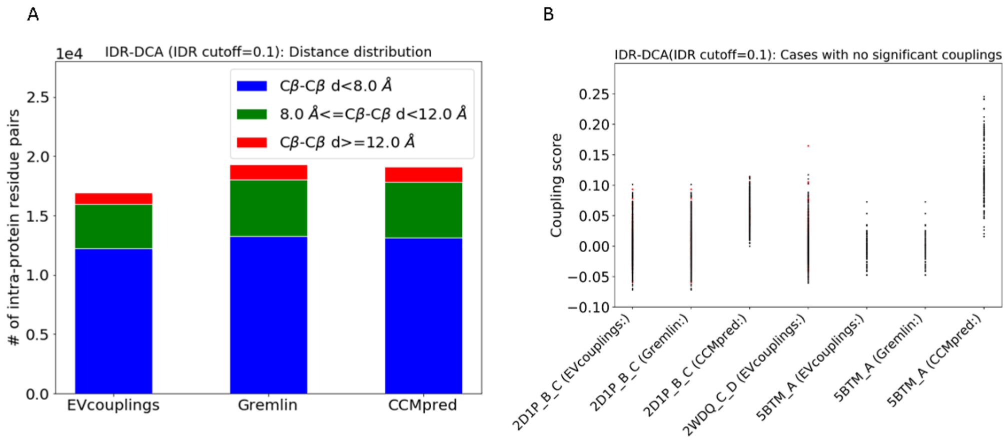


Figure S1. (A) The proportions of the intra-protein residue pairs selected by IDR-DCA (0.1 as the IDR cutoff) from the monomeric protein dataset with Cβ-Cβ distance smaller than 8Å, between 8Å and 12Å, and larger than 12 Å. (B) The coupling score scattering plot for the seven cases in which no evolutionary coupled residue pairs are selected by IDR-DCA (0.1 as the IDR cutoff), in which the contacting residue pairs are colored red, and the non-contacting residue pairs are colored black.

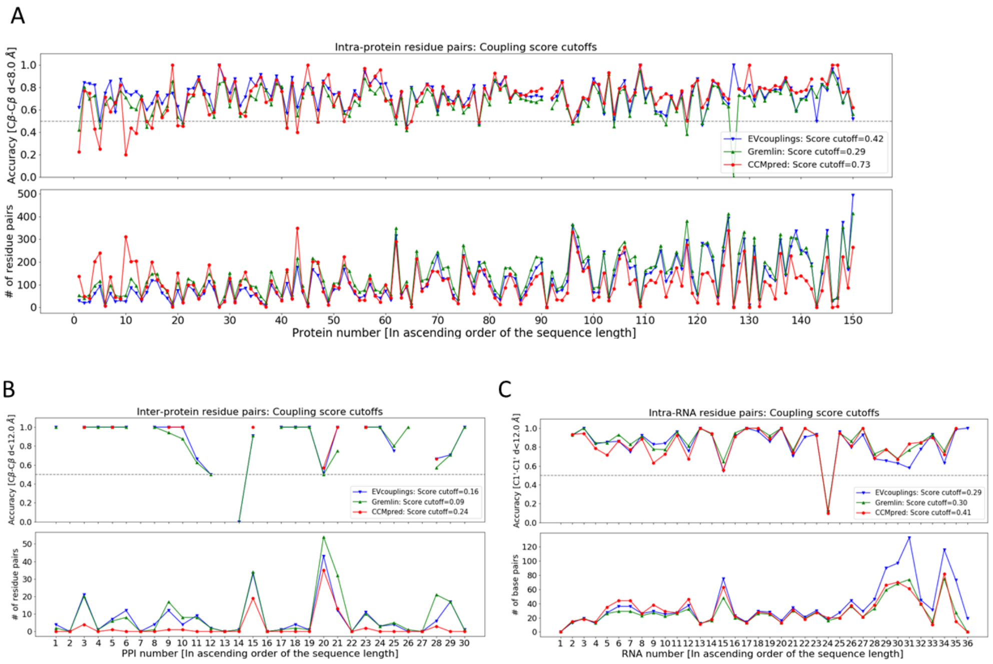


Figure S2. (A)~(C) The accuracies and the numbers of the residue pairs selected by applying the coupling score cutoffs for each case in the three datasets: (A) The monomeric protein dataset; (B) The protein-protein interaction dataset; (C) The monomeric RNA dataset. For each case, we applied EVcouplings, Gremlin and CCMpred to perform the DCA respectively. In the case that no residue pair is selected, the corresponding accuracy is not shown. For each dataset, the cases (proteins, protein-protein interactions, RNAs) are ordered ascendingly in the plot according to their sequence lengths.

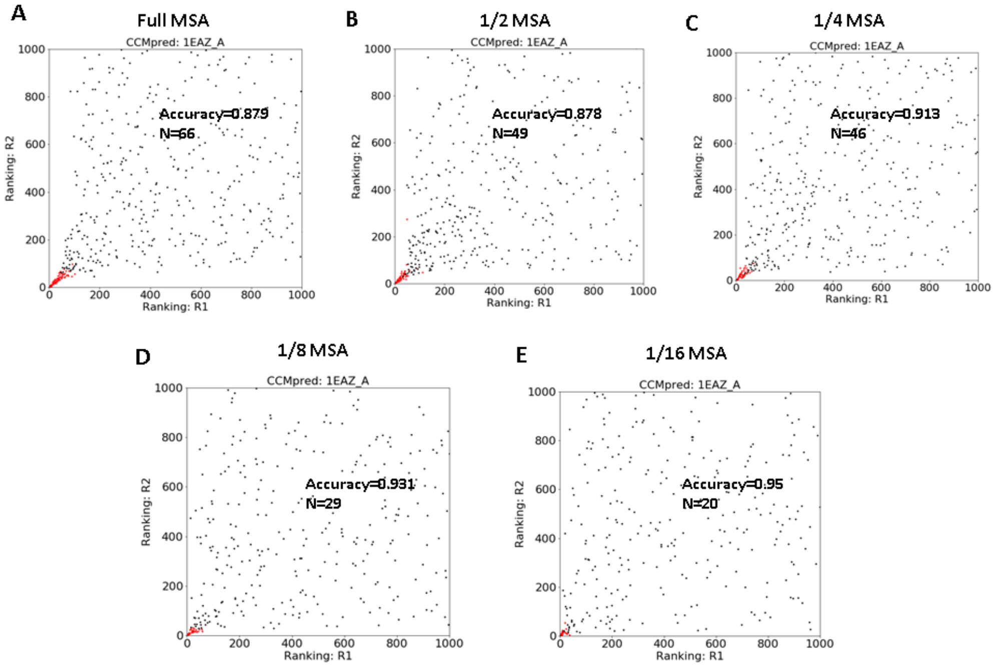


Figure S3. (A)-(E) An example of selecting evolutionary coupled residue pairs by applying IDR-DCA on the MSAs with different levels of downsampling: (A) Full MSA; (B) 1/2 MSA; (C) 1/4 MSA; (D) 1/8 MSA; (E) 1/16 MSA. In each ranking plot, the selected residue pairs are colored red, and the number and accuracy of the selected residue pairs are also shown.

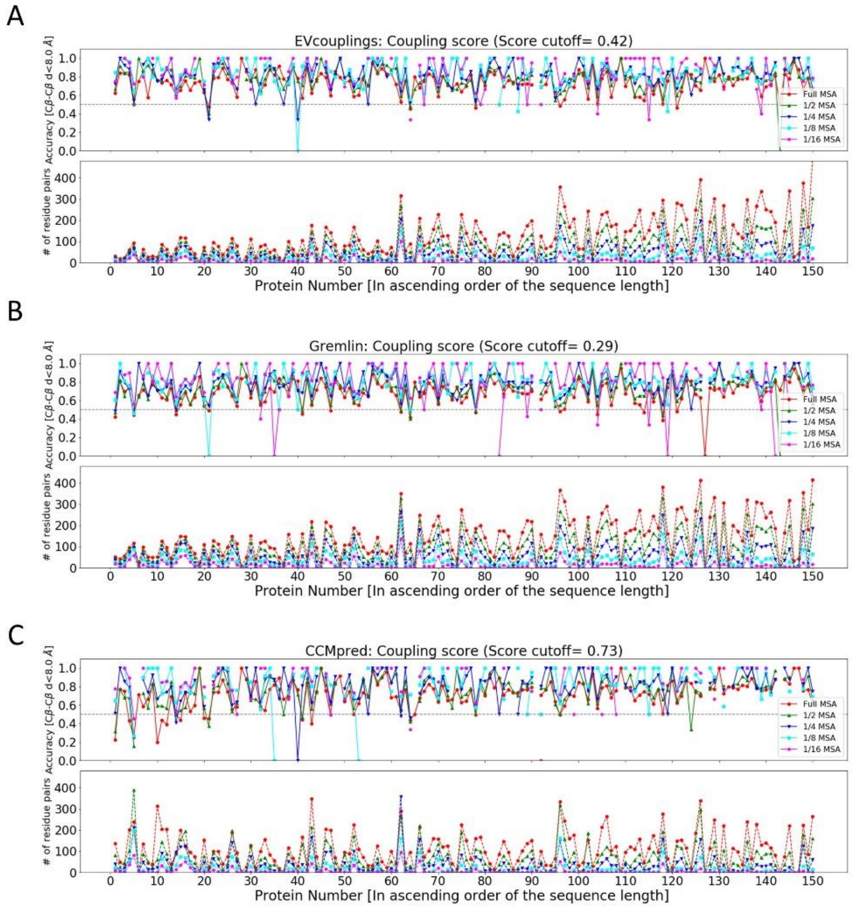


Figure S4. (A)-(C) The accuracies and the numbers of the residue pairs selected by applying the coupling score cutoffs on the MSAs with different levels of downsampling for each protein in the monomeric protein dataset, in which the DCA were performed with the application of the three DCA tools: (A) EVcouplings; (B) Gremlin; (C) CCMpred. In the case that no residue pair is selected, the corresponding accuracy is not shown. The proteins are ordered ascendingly according to their sequence lengths.

Table S1. The performances of RNA secondary structure prediction over 26 RNAs with the application of different types of prior base pair constraints

| PDB ID | Length | RNAFold (no constraints) | IDR-DCA | top L/5 | Coupling score |
|---|---|---|---|---|---|
| | | MCC | MCC | MCC | MCC |
| 4ENC_A | 52 | 0.44 | 0.48 | 0.48 | 0.48 |
| 5LYS_A | 56 | 1 | 1 | 1 | 1 |
| 4XWF_A | 64 | 0.75 | 1 | 1 | 1 |
| 4TZX_X | 71 | 0.98 | 0.98 | 0.98 | 0.98 |
| 3Q3Z_A | 75 | 0.28 | 0.97 | 0.97 | 0.97 |
| 1EHZ_A | 76 | 0.98 | 0.98 | 0.98 | 0.98 |
| 3D2G_A | 77 | 0.63 | 0.93 | 0.93 | 0.93 |
| 3CW5_A | 77 | 1 | 1 | 1 | 1 |
| 4YAZ_A | 84 | 0.63 | 0.9 | 0.79 | 0.9 |
| 3RG5_A | 86 | 1 | 1 | 1 | 1 |
| 3OX0_A | 86 | 0.94 | 0.94 | 0.94 | 0.94 |
| 4LVV_A | 89 | 0.74 | 0.91 | 0.83 | 0.91 |
| 4RUM_A | 91 | 0.97 | 0.97 | 0.97 | 0.97 |
| 3GX5_A | 94 | 0.93 | 0.93 | 0.93 | 0.93 |
| 4L81_A | 96 | 0.93 | 0.79 | 0.79 | 0.79 |
| 3SUH_X | 101 | 0.69 | 0.8 | 0.72 | 0.8 |
| 4WFL_A | 106 | 0.38 | 1 | 0.87 | 1 |
| 6DNR_A | 107 | 0.81 | 0.81 | 0.81 | 0.81 |
| 1NBS_A | 120 | 0.5 | 0.78 | 0.69 | 0.69 |
| 1U9S_A | 155 | 0.8 | 0.89 | 0.8 | 0.89 |
| 1GID_A | 158 | 0.84 | 0.84 | 0.84 | 0.81 |
| 3PDR_A | 161 | 0.78 | 0.78 | 0.78 | 0.78 |
| 3DIL_A | 173 | 0.9 | 0.98 | 0.94 | 0.98 |
| 1Y0Q_A | 229 | 0.54 | 0.88 | 0.95 | 0.88 |
| 2A64_A | 298 | 0.6 | 0.81 | 0.81 | 0.76 |
| 1FFZ_A | 496 | 0.59 | 0.83 | 0.84 | 0.59 |
| **Mean1** | **126** | **0.75** | **0.89** | **0.87** | **0.88** |
| **Mean2(length>100)** | **191** | **0.68** | **0.85** | **0.82** | **0.82** |

Data S1. The raw data for Figure 3.

Data S2. The raw data for Figure S2.
